# African *Campylobacter jejuni* genomes reveal globally connected population structure and regionally variable resistance, virulence and mobilome profiles

**DOI:** 10.64898/2026.07.27.741116

**Authors:** Abdalah Makaranga, Samweli Bahati, Eliezer Mwakalapa, Henry Mung’ong’o, Albino Kalolo, Claus Thomas, Daud Kassam, Pilirani Chisembe, Keigo Shibayama, Reuben S. Maghembe

## Abstract

*Campylobacter jejuni* is a leading foodborne cause of gastroenteritis, but genomic surveillance remains uneven across Africa. A recent East Africa study combined whole-genome sequencing and antimicrobial susceptibility testing for *Campylobacter* isolates from humans with diarrhea in Kenya and poultry in Tanzania, showing high sequence-type diversity and substantially higher multidrug resistance in poultry. We extended this regional evidence by analyzing 1,013 publicly available *C. jejuni* genomes, including 718 African and 295 non-African comparator genomes, with standardized assembly, genotyping, phylogenomics, pangenome reconstruction, antimicrobial resistance, virulence, and mobile-element profiling. African genomes were geographically concentrated but genetically diverse, included globally distributed and regionally enriched lineages, and showed an open pangenome dominated by low-frequency gene families. Resistance and virulence determinants were unevenly distributed by region and lineage. These findings place African *C. jejuni* diversity within a global evolutionary framework and support expanded, integrated One Health genomic surveillance.

**Data summary:** All genome sequence data analysed in this study were retrieved from publicly accessible repositories, including the National Center for Biotechnology Information Sequence Read Archive and Assembly resources and corresponding records available through the International Nucleotide Sequence Database Collaboration where applicable. Accession identifiers, BioSample records, run accessions, country metadata and host/source information for all analysed genomes are provided in the combined Supplementary Data workbook. No new sequence data were generated. The analysis used publicly available data generated by other investigators, and the original data-generating studies are cited where appropriate. Derived analytical outputs supporting the findings are included in the manuscript and Supplementary Information.

**Impact statement:** Genomic surveillance of *Campylobacter jejuni* remains uneven globally, and African data are still underrepresented in many comparative analyses. This study brings together publicly available African and non-African *C. jejuni* genomes in a single standardized comparative framework, linking population structure, pangenome composition, antimicrobial-resistance determinants, virulence-associated loci and mobile-element profiles. The work shows that African *C. jejuni* diversity is not peripheral to the global population: African genomes include globally distributed sequence types, regionally enriched lineages and a large accessory-gene repertoire. By separating genomic surveillance signals from population-representative prevalence claims, the study provides a cautious framework for interpreting public genome collections from settings with unequal sampling. The findings support broader One Health genomic surveillance, improved metadata completeness and geographically balanced sequencing to better understand foodborne transmission, resistance evolution and lineage diversification in this important zoonotic pathogen.

## Introduction

*Campylobacter jejuni* is one of the leading causes of bacterial gastroenteritis worldwide, responsible for a substantial burden of diarrhoeal disease across both high-income and low-and middle-income countries [1–3]. In the WHO global foodborne-disease estimates, diarrhoeal agents accounted for most foodborne illnesses; *Campylobacter* spp. Caused approximately 96 million foodborne illnesses in 2010 and ranked among the most frequent enteric hazards, alongside norovirus and major bacterial pathogens such as non-typhoidal Salmonella enterica, enteropathogenic and enterotoxigenic *Escherichia coli*, and *Shigella* spp. [4]. Transmission occurs primarily through contaminated food products, especially poultry, as well as through environmental and animal reservoirs, reflecting its broad ecological niche and zoonotic potential [1,5,6]. Despite its global importance, the epidemiology and evolutionary dynamics of *C. jejuni* remain poorly characterised in many African settings.

Advances in whole-genome sequencing (WGS) have transformed our understanding of bacterial population structure, enabling high-resolution analysis of lineage diversification, transmission pathways, and functional adaptation [7–9]. In *C. jejuni,* genomic studies have revealed a complex population structure by both globally disseminated generalist lineages and locally adapted strains associated with specific hosts or ecological niches (1, 2, 6). However, much of this knowledge is derived from datasets dominated by isolates from Europe, North America, and parts of Asia [12–15], creating a significant geographic bias that limits global inference.

Africa represents a critical gap in *C. jejuni* genomic surveillance. The continent is characterised by diverse ecological systems, varying agricultural practices, and differing patterns of antimicrobial use, all of which can influence pathogen evolution and transmission. Yet, publicly available genomic data from Africa remain sparse and unevenly distributed, constraining our ability to understand the contribution of African populations to global *C. jejuni* diversity and the emergence of region-specific lineages or resistance profiles [16–18]. Addressing this gap is essential for building a truly global framework of *C. jejuni* evolution and for informing context-specific public health interventions.

At the genomic level, *C. jejuni* exhibits a highly dynamic genome architecture, with an open pangenome characterized by extensive accessory gene variation [12,15,19]. This genomic plasticity underpins the organism’s ability to adapt to diverse hosts and environments, but also complicates efforts to define stable genetic markers of virulence and antimicrobial resistance (AMR). While certain resistance mechanisms, such as the *cmeABC* efflux system and fluoroquinolone-associated mutations, are widely conserved, the distribution of other resistance determinants varies across populations and geographic regions [18,20]. Similarly, although core virulence factors are broadly conserved, variation in accessory virulence-associated loci may contribute to differences in pathogenic potential and host adaptation [12,19]. In parallel, the role of mobile genetic elements in *C. jejuni* evolution appears to differ from that in many other bacterial pathogens. While horizontal gene transfer contributes to genetic diversity, plasmid-mediated dissemination of resistance and virulence factors is less prominent, suggesting that genome plasticity and recombination may play a more central role in shaping population structure [9,21]. Understanding how these processes interact at a global scale, and whether they vary across regions, remains a key challenge.

The 2024 East Africa Emerging Infectious Diseases study by French and colleagues provides the closest regional comparator for the present work. That study analyzed 178 *Campylobacter* isolates from humans with diarrhea in Kenya and poultry in Tanzania and showed high sequence-type diversity, shared sequence types between humans and poultry, and a markedly higher prevalence of multidrug resistance in poultry isolates than in human isolates [22]. Building on that isolate-based East Africa evidence, we performed a broader comparative genomic analysis of 1,013 *C. jejuni* genomes, including 718 African genomes and 295 non-African comparators. By integrating phylogenomics, pangenome analysis, antimicrobial resistance profiling, virulence-factor screening, and mobile-element analysis, we aimed to place African *C. jejuni* diversity within a global evolutionary and surveillance framework.

## Methods

### Dataset description and study design

#### African genome landscape

Publicly available whole-genome sequencing (WGS) data for *C. jejuni* were retrieved from international sequence repositories up to 30 September, 2025. These included the National Center for Biotechnology Information (NCBI) Sequence Read Archive/Assembly resources and records mirrored through the International Nucleotide Sequence Database Collaboration, including the European Nucleotide Archive (ENA) and DNA Data Bank of Japan (DDBJ), where relevant accession metadata were available. Repository-derived accession identifiers, BioSample records, run accessions, country information, host/source metadata, and the variables extracted for analysis are provided in Supplementary Data 1. Metadata were curated to standardize the country of origin and host source. All genomes originating from African countries were retained at this stage, without filtering by sequencing strategy or data quality, to describe the continental availability and sampling landscape of *C. jejuni* WGS data.

#### Definition of the analyzed dataset

To enable comparative genomic analyses using standardized sequencing data, the African genome collection was filtered to retain only isolates generated using paired-end Illumina sequencing with publicly available raw reads. This resulted in a subset of 718 African *C. jejuni* genomes, which were used as representative African isolates for all downstream analyses.

To provide a global comparative context, a set of 295 non-African paired-end Illumina genomes was assembled from countries selected based on (i) availability of raw paired-end sequencing data suitable for standardized downstream analysis and (ii) documented epidemiological relevance of *C. jejuni* across human, animal, and environmental reservoirs [23–28]. The comparator dataset includes genomes from Japan, China, Brazil, Spain, Peru, Canada, Norway, Viet Nam, the United States, the United Kingdom, Sweden, and India, representing diverse geographic regions, surveillance capacities, and transmission settings. This approach enables placement of African isolates within a broader global population structure while ensuring data quality, comparability, and representation of epidemiologically relevant lineages. Detailed isolate-level metadata for both African and non-African genomes, including country of origin, host source, year of isolation, sequencing platform, and accession identifiers, are provided in Supplementary Data 1.

#### Read quality control and genome assembly

Raw paired-end Illumina sequencing reads from all 1,013 *C. jejuni* genomes (718 African and 295 non-African) were processed using fastp v1.0.1 [29]. The same quality-control workflow was applied to all samples to avoid technical bias between regions. No datasets were excluded at this stage, as all samples satisfied minimum quality requirements for short-read bacterial genome assembly. Quality-controlled reads were assembled de novo using SPAdes version 4.2.0 [30], in short-read mode for all 1,013 genomes. Contigs shorter than 500 bp were removed to minimize assembly artefacts. Assembly quality was assessed using standard metrics, including total assembly length, number of contigs, largest contig size, N50, GC content, and misassembly statistics. Assemblies were retained if their total length fell within the expected *C. jejuni* genome size range (approximately 1.5–2.1 Mb).

### Species confirmation and basic genotyping

Species confirmation and baseline genotyping were performed using the Pathogenwatch platform [31]. For each assembled genome, Pathogenwatch was used to verify that the genome was consistent with *C. jejuni* based on its internal species assignment framework, and to generate multilocus sequence typing (MLST) outputs. MLST sequence types (STs) were extracted directly from Pathogenwatch outputs and merged into the master metadata table for downstream analyses.

#### Genome annotation and pangenome analysis

Genome annotation was performed for all final included *Campylobacter jejuni* assemblies using Bakta v1.11 [32], applying default parameters to generate standardized gene predictions across the dataset. Pangenome analysis was conducted using PPanGGOLiN v2.2.2 [33], on the complete set of annotated genomes. Gene families were inferred based on sequence similarity and partitioned into pangenome categories using PPanGGOLiN’s graph-based model. Gene families were classified into core, soft-core, shell, and cloud partitions according to their frequency across genomes. Core genes were defined as those present in at least 99% of genomes, soft-core genes as those present in 95–99% of genomes, shell genes as those present in 15–95% of genomes, and cloud genes as those present in fewer than 15% of genomes.

### Phylogenomic reconstruction and population structure

Roary v3.11.2 [34], was used to generate the core gene alignment. Maximum-likelihood phylogenetic inference was performed using IQ-TREE2 v2.4.0 [35], on the Roary-derived core alignment. Major clades were defined as well-supported monophyletic groups corresponding to the deepest bifurcations in the tree topology. Sublineages were delineated as nested monophyletic clusters within each major clade, identified based on consistent branching structure and relative branch length separation. Clonal complexes were assigned using the PubMLST *Campylobacter* scheme (http://pubmlst.org/Campylobacter) by mapping each genome’s MLST sequence type (ST) and allele profiles to the corresponding CC definition. Visualization was done by using iTOL v7 [36].

### Antimicrobial resistance and virulence factor profiling

Antimicrobial resistance (AMR) determinants were identified across all 1,013 confirmed genomes using two complementary approaches. First, assembled genomes were screened with ABRicate against curated resistance gene databases. In parallel, resistance profiling was performed using the Resistance Gene Identifier (RGI) (v6.0.5) with the Comprehensive Antibiotic Resistance Database (CARD) (v4.0.1) [37]. Default parameters were applied for both tools, and only high-confidence hits based on database-specific scoring criteria were retained for downstream comparative analysis. Virulence-associated determinants were identified across all 1,013 confirmed genomes using ABRicate. Assembled genomes were screened against the Virulence Factor Database (VFDB) [37] to detect known virulence-associated loci. Default parameters were applied, and hits were filtered based on standard sequence identity and coverage thresholds to retain high-confidence matches. The resulting virulence profiles were used for downstream comparative analyses across isolates.

### Plasmid and mobile genetic element analysis

Plasmid sequences and associated mobile genetic elements were identified using MOB-suite. Genome assemblies were analyzed to reconstruct putative plasmids, assign replicon types, and infer plasmid mobility based on relaxase typing, mating pair formation (MPF) systems, and origin of transfer (oriT) sequences.

### Statistical analysis

Statistical analyses were aligned with the major analytical objectives of the study. The African genome landscape was summarized descriptively by country, host/source, sequencing metadata, and genome availability to quantify geographic and sampling imbalance. Assembly quality metrics were summarized using counts, ranges, means, and proportions to assess whether genomes met expected *C. jejuni* assembly characteristics. Population diversity was assessed by tabulating MLST sequence types, clonal complexes, host/source categories, and country distributions, followed by comparison of shared and region-specific STs between African and non-African datasets. Pangenome structure was summarized by the number and proportion of gene families assigned to core, soft-core, shell, and cloud partitions. Phylogenomic structure was interpreted from the maximum-likelihood topology by defining major clades and sublineages and then summarizing their ST, clonal-complex, host/source, and country composition. For antimicrobial resistance, virulence, plasmid, and mobile genetic element analyses, prevalence of each determinant was calculated as the proportion of positive isolates (n/N) in African and non-African datasets. Odds ratios (ORs) were estimated using the Haldane–Anscombe correction to account for zero counts and ensure stable estimates [38]. 95% confidence intervals (CIs) were calculated on the log scale and back-transformed to the OR scale. Statistical significance of differences in prevalence was assessed using two-sided Fisher’s exact test, with p < 0.05 considered significant.

### Interpretive scope of comparative analyses

Because the comparator collection was assembled to provide genomic context rather than population-representative prevalence estimates, regional comparisons were interpreted as differences within the analysed public-genome collection. Determinant-level comparisons were treated as exploratory and interpreted alongside lineage composition and unequal repository sampling.

## Results

### Geographic landscape of African *C. jejuni* genomes

A total of 784 Campylobacter jejuni genomes originating from Africa were identified as of 30 September, 2025 (Fig. 1). Genome availability was dominated by Ethiopia (n = 326), followed by Kenya (n = 105), Tanzania (n = 79), Uganda (n = 71), Egypt (n = 73), and Botswana (n = 64). Additional contributions were observed from Mali (n = 34) and Gambia (n = 28), while Burkina Faso (n = 2) and South Africa (n = 2) were minimally represented. Large regions of Central and North Africa lacked publicly available genomes.

**Fig. 1.**
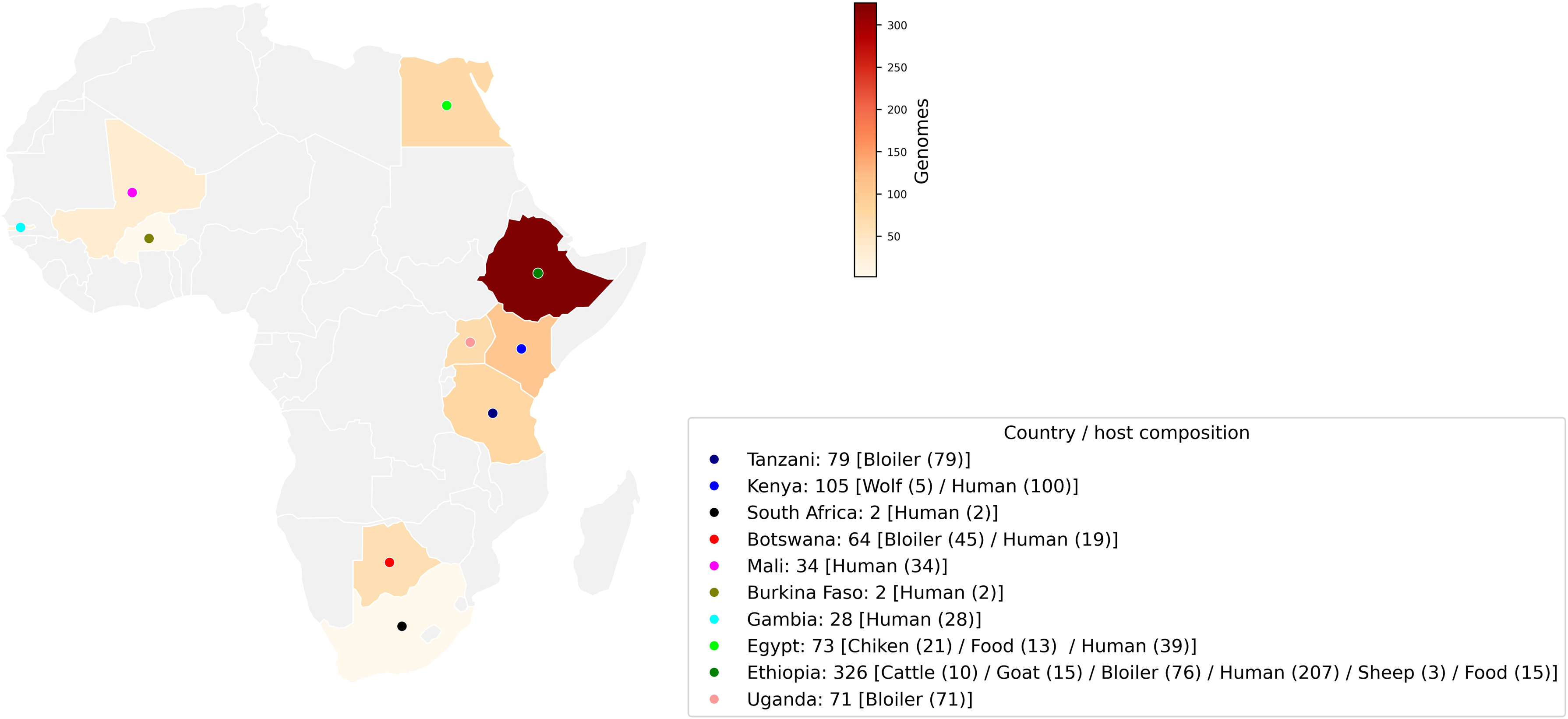
African Campylobacter jejuni whole-genome sequencing landscape as of 27 August 2025. The choropleth map shows the number of publicly available C. jejuni genomes per African country (n = 784 total). Countries without available genomes are shown in grey. The distribution highlights pronounced geographic imbalance, with the majority of genomes originating from a small number of countries, particularly in East Africa.

Host-associated metadata indicated that African *C. jejuni* genomes were predominantly derived from human clinical samples and poultry (broiler) sources, with additional contributions from livestock and food-associated samples in a small number of countries. Notably, host composition varied markedly between countries, reflecting differences in local surveillance priorities and study designs rather than continent-wide sampling uniformity. More details in Supplementary Data 1.

### Final dataset and assembly quality

After uniform processing of the 1,013 paired-end datasets, mean post-filtering Q20 and Q30 values were 98.0% and 95.4%, respectively, and mean GC content remained consistent with *C. jejuni.* All assemblies fell within the expected genome-size range (mean, approximately 1.85 Mb), but contiguity varied substantially (14–10,424 contigs; N50 approximately 8.7 kb to >1.08 Mb) (Supplementary Data 1).

### Species confirmation and population diversity

All included assemblies were Campylobacter jejuni. Multilocus sequence typing (MLST) assignments were successfully obtained for the majority of genomes, yielding a diverse distribution of sequence types (STs) across African and non-African isolates (Fig. 2). Detailed isolate-level ST assignments, host sources, and country information are provided in Supplementary Data 1.

**Fig. 2.**
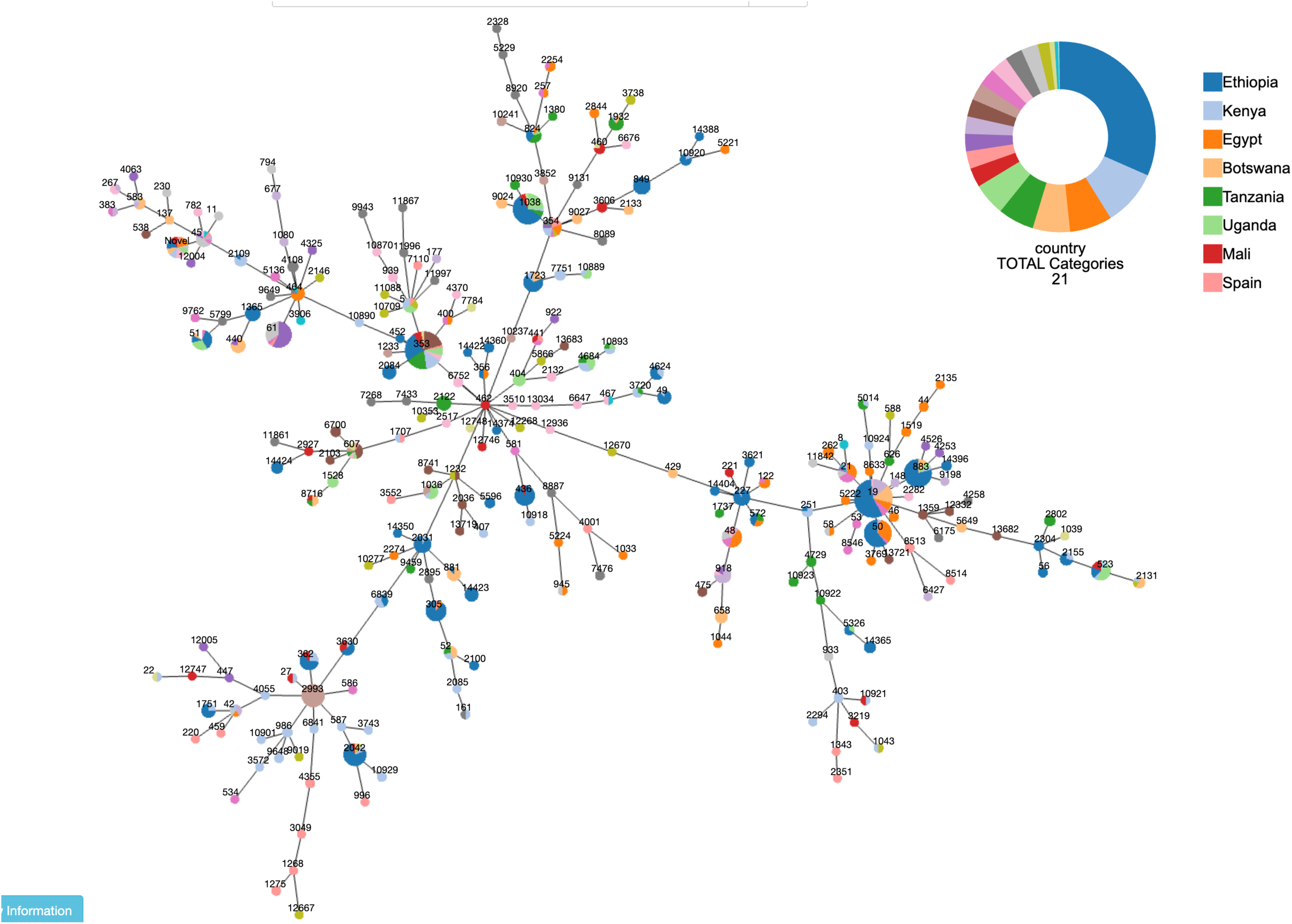
Multilocus sequence type (ST) distribution of *Campylobacter jejuni* isolates based on MLST analysis. The network visualization shows the distribution and relationships of multilocus sequence types (STs) identified among the analyzed *C. jejuni* genomes. Each node represents an ST, with node size proportional to the number of isolates assigned to that ST. Node colors indicate country of origin, as shown in the accompanying legend. Edges connect STs sharing allelic relationships within the MLST scheme. The donut chart summarizes the proportional contribution of countries to the overall ST dataset, highlighting the relative representation of isolates from different geographic origins.

African genomes showed extensive sequence-type diversity, led by ST-19 (n = 35), ST-1038 (n = 33), and ST-353 (n = 32), each occurring across multiple countries and host sources; Pathogenwatch reported 14 novel STs. Non-African comparators were also diverse, led by ST-61 (n = 26) and ST-2993 (n = 21). Several STs, including ST-19, ST-353, ST-50, ST-354, ST-48, ST-305, ST-440, and ST-464, occurred in both datasets, indicating globally distributed lineages alongside regional diversity (Supplementary Data 1).

### Pangenome composition

Pangenome reconstruction identified 48,046 gene families, including 889 core, 288 soft-core, 929 shell, and 45,940 cloud families. The dominance of low-frequency cloud families, together with the U-shaped gene-frequency distribution, indicates an open pangenome with extensive accessory-genome variation (Fig. 3).

**Fig. 3.**
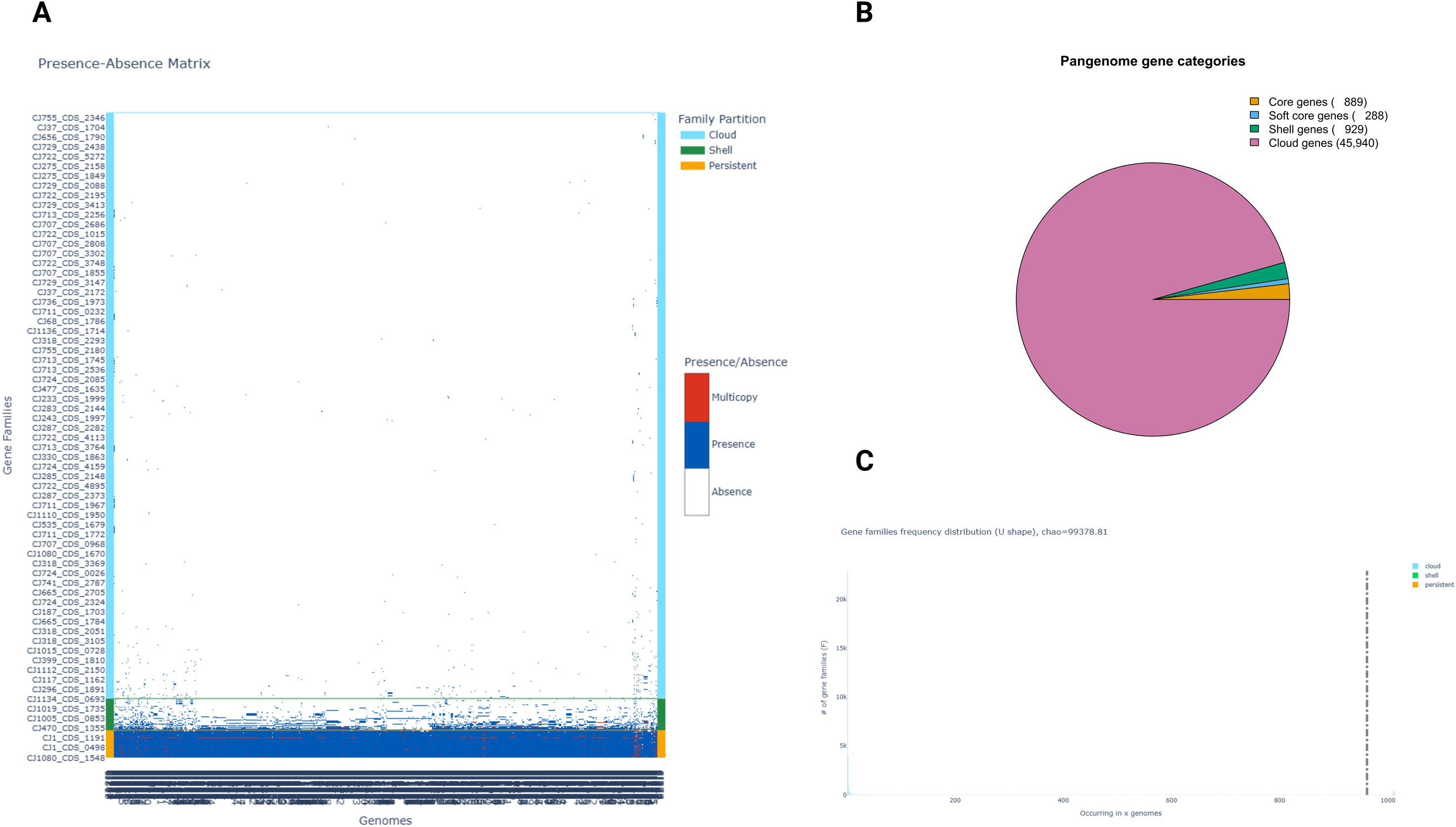
Pangenome structure and gene family distribution of *Campylobacter jejuni* isolates. **(A)** Presence–absence matrix of gene families across all analyzed genomes, showing the distribution of gene families categorized into persistent (core), shell, and cloud partitions. Each row represents a gene family, and each column represents a genome, with color-coded signals indicating gene presence, absence, or multicopy occurrence. The matrix highlights the high prevalence of core genes and extensive variability in accessory gene content across isolates. **(B)** Proportional distribution of pangenome gene categories, illustrating a highly open pangenome dominated by cloud genes (45,940 families), with smaller contributions from shell (929), soft-core (288), and core genes (889). **(C)** Frequency distribution of gene families across genomes, displaying a characteristic U-shaped pattern, where most gene families are either rare (cloud genes) or highly conserved (core genes), consistent with an open pangenome architecture.

### Phylogenomic structure and evolutionary diversification

Core-gene phylogeny of 1,002 genomes resolved two major clades, MCI (n = 762) and MCII (n = 240) (Fig. 4). MCI included a CC-21-dominated sublineage spanning human and chicken sources and an East Africa-centred CC-353/CC-354 radiation. MCII comprised smaller CC-45- and CC-49/CC-61-centred sublineages and a human-associated lineage containing ST-2993, ST-2042, and ST-362. Host and country distributions across clades indicate both intercontinental lineage sharing and regional enrichment.

**Fig. 4.**
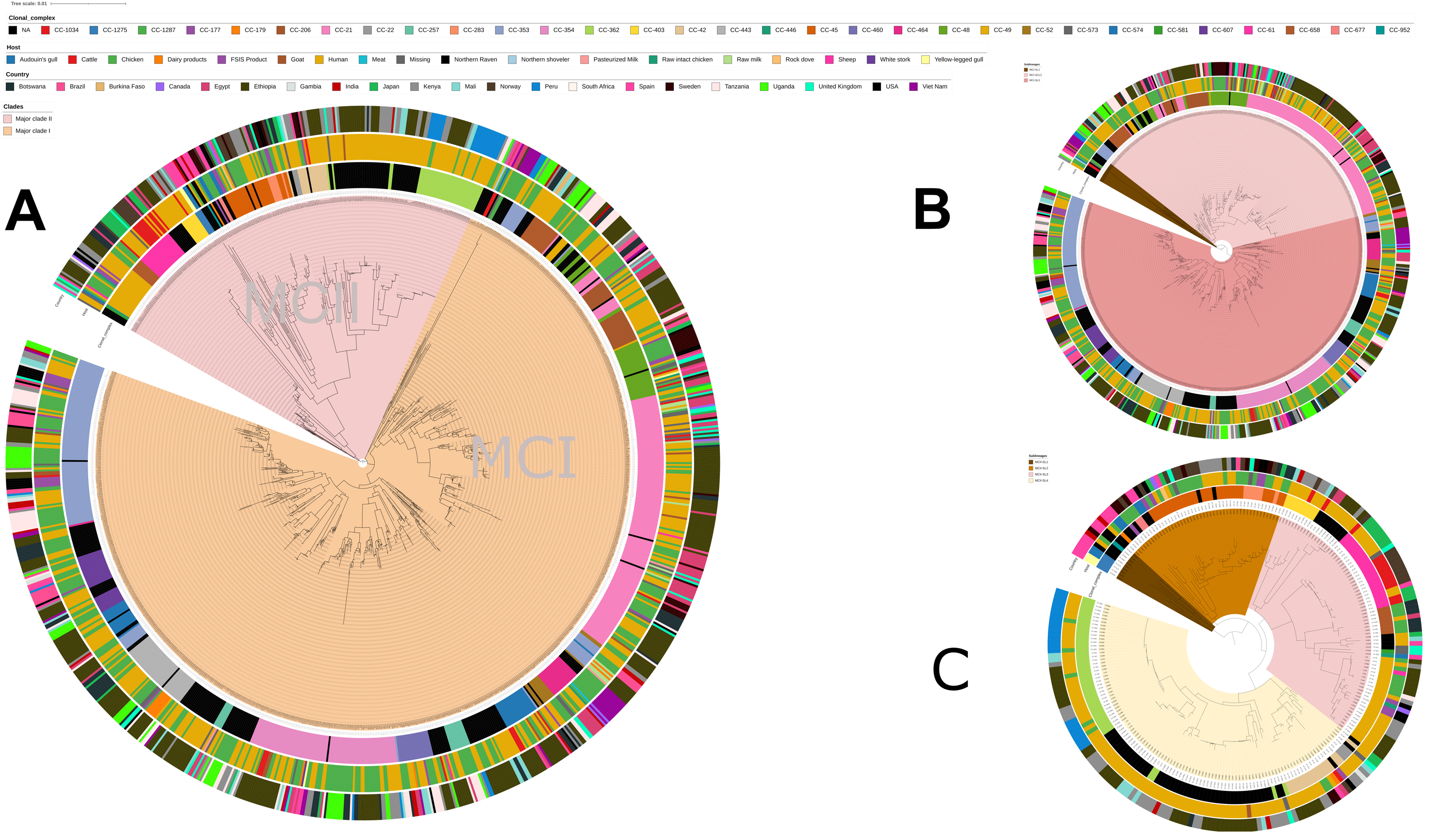
Phylogenomic structure and clade stratification of *Campylobacter jejuni* isolates. **(A)** Maximum-likelihood phylogeny revealing two major phylogenetic clades (MCI and MCII). Concentric annotation rings represent clonal complex (CC), host/source, country of origin, and clade assignment, illustrating the global diversity and extensive mixing of isolates across geographic regions and ecological niches. **(B)** Subtree corresponding to Major Clade I (MCI), highlighting its internal phylogenetic structure and lineage diversification. The distribution of clonal complexes, hosts, and countries indicates the presence of both widely distributed and lineage-specific clusters within this clade. **(C)** Subtree corresponding to Major Clade II (MCI), showing a distinct phylogenetic organization. The clustering patterns suggest differences in lineage composition and potential host or geographic associations within this clade.

#### Antimicrobial resistance determinants

**T**he AMR matrix comprised 1,013 genomes and showed a compact but uneven resistance repertoire dominated by four highly prevalent determinants: *cmeA, cmeC*, and *cmeR* were each present in 99.7% of genomes, and *cmeB* in 94.7%, followed by *OXA-450* (63.2%), *gyrA*_T86I (36.0%), and *tetO* (33.6%), whereas all remaining markers occurred at ≤4.7% (Fig. 5).

**Fig. 5.**
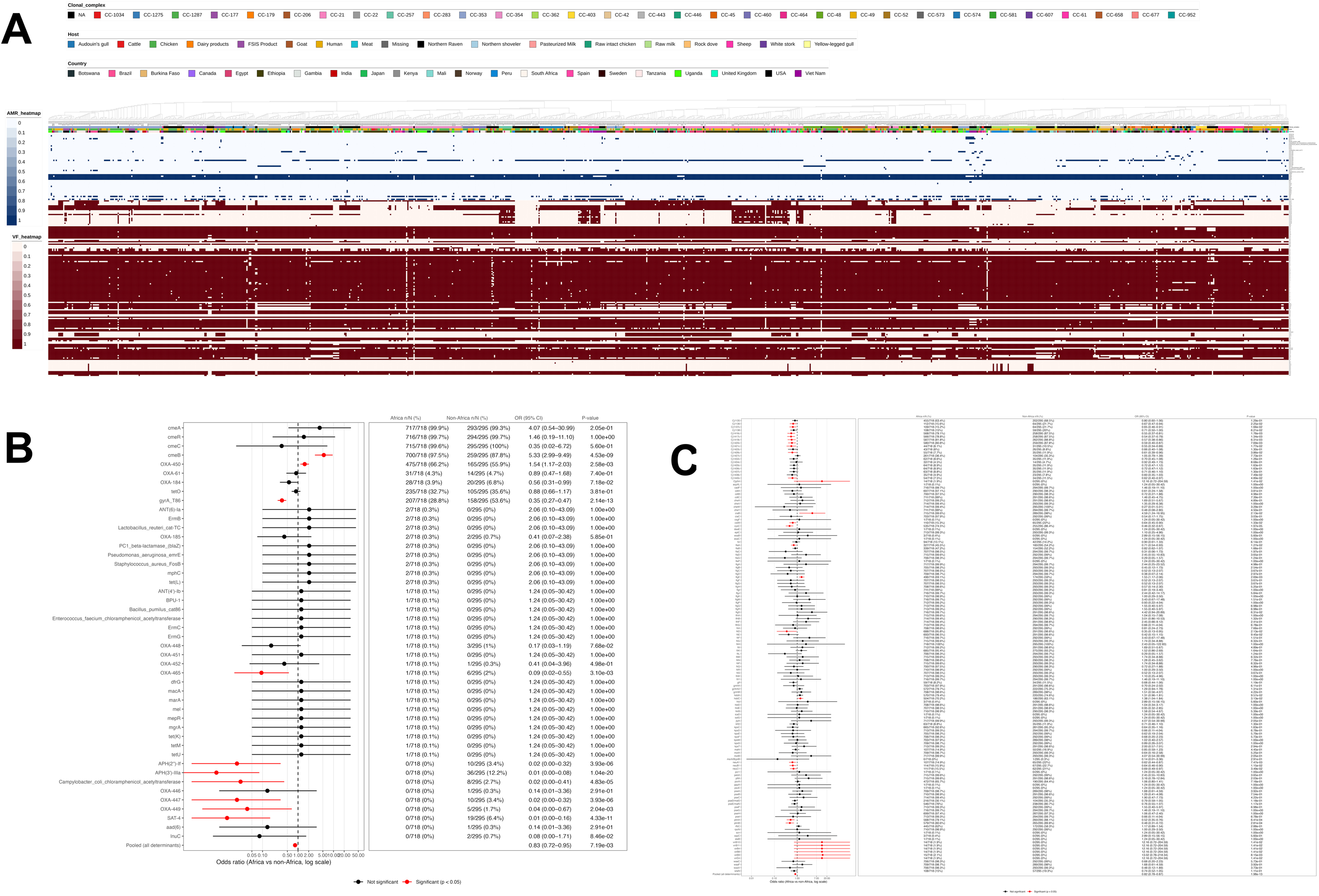
Integrated resistome and virulome profiling with comparative statistical analysis of *Campylobacter jejuni* isolates. **(A)** Heatmap showing the distribution of antimicrobial resistance (AMR) determinants (blue scale) and virulence factors (VF; red scale) across all genomes, with isolates ordered and annotated by clonal complex (CC), host/source, and country of origin. The AMR profile reveals a relatively sparse and heterogeneous distribution of resistance determinants, whereas virulence factors exhibit high prevalence and conservation across isolates, reflecting core pathogenic traits. **(B)** Forest plot summarizing the comparative prevalence of AMR determinants between African and non-African isolates. Odds ratios (log scale) with 95% confidence intervals are shown for each determinant, with significant differences (p < 0.05) highlighted in red, indicating region-specific enrichment or depletion of selected resistance genes. **(C)** Forest plot of virulence factor determinants comparing African and non-African isolates, demonstrating largely conserved virulence profiles with limited but notable differences in specific loci. The narrow confidence intervals around most estimates indicate stable prevalence across populations, with only a subset of determinants showing statistically significant regional variation.

Among individual AMR markers, *cmeB* (97.5% vs 87.8%; OR = 5.33, 95% CI 2.99–9.49) and OXA-450 (66.2% vs 55.9%; OR = 1.54, 95% CI 1.17–2.03) were enriched among African genomes in this collection, whereas gyrA_T86I was more frequent among non-African comparators (28.8% vs 53.6%; OR = 0.35, 95% CI 0.27–0.47) (Fig. 5). Several low-frequency determinants occurred only or predominantly among non-African comparator genomes; full determinant-level comparisons are provided in Supplementary Data 2.

### Virulence factor distribution

Virulence screening detected a widely conserved core repertoire, including near-universal flagellar and adhesion-associated loci. Among variable loci, ptmA/ptmB, maf4, and pseE/maf5 were more frequent among African genomes, whereas porA, Cj1135, rfbC, and rare virB-associated loci were more frequent among non-African comparators (Fig. 5; Supplementary Data 2). These findings indicate a conserved virulence backbone overlaid by lineage- and region-associated variation.

### Plasmid architecture and mobile genetic element distribution

Mobilome profiles were sparse and broadly similar across datasets. Replicons, relaxases, and mating-pair-formation systems occurred in 18.0%, 24.2%, and 23.7% of African genomes, respectively, and in 15.6%, 26.8%, and 26.1% of non-African comparator genomes. Most genomes were classified as non-mobilizable, and insertion-sequence burden was low, supporting limited large-plasmid diversity in this collection (Supplementary Data 3 and Supplementary Data 4).

## Discussion

In this large-scale genomic analysis of 1,013 *C. jejuni* isolates, we provide a comprehensive assessment of African population structure within a global framework, integrating phylogenomics, pangenome architecture, antimicrobial resistance (AMR), virulence determinants, and mobile genetic elements. Our findings show that *C. jejuni* in Africa is embedded within globally distributed lineages but exhibits distinct regional structuring, characterised by extensive genomic diversity, a highly open pangenome, and a conserved functional backbone overlaid by geographically differentiated resistance and virulence profiles.

Our results complement and extend the 2024 East Africa study by French and colleagues, which focused on contemporaneous WGS and phenotypic antimicrobial susceptibility testing of human and poultry *Campylobacter* isolates from Kenya and Tanzania [18]. Whereas that study was particularly strong for host-linked interpretation and poultry-associated multidrug resistance, the present analysis broadens the question to a continental and global scale. The two studies therefore address different but connected surveillance needs: isolate-linked One Health evidence in East Africa and large-scale genomic contextualization of African *C. jejuni* diversity within the global population.

A striking feature of the dataset is the pronounced geographic imbalance in genome availability across Africa (Fig. 1), with sequencing efforts concentrated in a limited number of countries, particularly in East Africa. This uneven representation likely reflects disparities in sequencing capacity and surveillance infrastructure rather than true epidemiological distribution. Similar limitations in global pathogen genomic surveillance have been highlighted in recent studies, which show that underrepresentation of low-resource regions constrains accurate inference of pathogen evolution and transmission dynamics [39,40]. Nevertheless, the available African dataset already captures substantial diversity, suggesting that current data represent only a fraction of the true genomic landscape.

Phylogenomic reconstruction resolved two deeply divergent clades (MCI and MCII), reflecting an early evolutionary split followed by extensive lineage-specific diversification (Fig. 4). Within these clades, we observe a dual evolutionary pattern comprising both clonal expansion and ongoing diversification. Major sublineages are structured around globally disseminated clonal complexes such as CC-21 and CC-353, which are widely recognised as host-generalist lineages with enhanced ecological fitness and transmission capacity [9–11]. The persistence and global distribution of these lineages across multiple host reservoirs in our dataset reinforce their central role in *C. jejuni* population structure. At the same time, the presence of numerous low-frequency and novel sequence types within African isolates shown in Fig. 2 indicates ongoing local diversification within this globally connected population. The coexistence of shared sequence types across continents (e.g., ST-19, ST-353, ST-50) highlights extensive intercontinental connectivity consistent with the zoonotic and foodborne transmission ecology of *C. jejuni* [10,41]. However, the simultaneous presence of regionally enriched lineages suggests that global dissemination is accompanied by geographically structured evolution, likely shaped by local ecological conditions, agricultural practices, and host interactions.

Pangenome analysis reveals a highly open genome dominated by a vast cloud gene repertoire (Fig. 3), with the majority of gene families occurring at low frequency. This architecture reflects continuous gene gain and loss and indicates strong evolutionary plasticity. In contrast, the relatively small core genome represents a conserved functional backbone required for essential biological processes. This U-shaped distribution is characteristic of bacterial species with broad ecological niches and is consistent with recent studies on bacterial pangenome dynamics, which demonstrate that accessory genome expansion underpins ecological adaptability and niche diversification [9,19]. In this context, much of the adaptive potential of *C. jejuni* likely resides within its accessory genome rather than its conserved core.

Despite this extensive genomic variability, the resistome is comparatively constrained and dominated by a small number of highly prevalent determinants (Fig. 5). The near-universal presence of the cmeABC efflux system and its regulator cmeR underscores the central role of intrinsic resistance mechanisms in *C. jejuni*, as also reported from other studies [20,42]. These systems provide baseline resistance across multiple antimicrobial classes and are highly conserved across lineages. In contrast, acquired resistance determinants are relatively sparse and unevenly distributed, indicating that resistance evolution is driven primarily by chromosomal mutations and modulation of intrinsic systems rather than extensive horizontal gene acquisition [19,42]. We identified significant regional differences in AMR profiles, with African genomes in this public-data collection showing lower overall detected determinant prevalence but enrichment of specific markers, including *cmeB* and *OXA-450*, while mutations such as *gyrA*_T86I and several aminoglycoside resistance genes were more prevalent in non-African genomes (Fig. 5). These findings are consistent with global analyses demonstrating that antimicrobial resistance patterns reflect regional antibiotic usage in both human and agricultural settings [20,41,43,44]. The absence of several resistance determinants among available African genomes suggests geographically structured occurrence, but requires confirmation in balanced surveillance datasets.

In contrast to the resistome, the virulence factor repertoire is highly conserved, with most determinants present in nearly all genomes (Fig. 5). This conservation suggests that core virulence functions are essential for *C. jejuni* biology and are maintained across diverse lineages and environments [8,42,44]. However, significant regional differences in a subset of loci, including enrichment of *ptmA/ptmB* and *maf*-associated genes in African isolates and higher prevalence of porA and virB-associated loci in non-African genomes, indicate fine-scale adaptation within this conserved framework. The absence of virB-associated genes in African isolates is particularly notable and suggests geographic structuring of specific virulence strategies, although the functional implications of this pattern require further investigation.

The mobilome analysis revealed a relatively sparse plasmid and mobile genetic element landscape, with most genomes lacking mobilizable elements and only a minority classified as conjugative. This contrasts with many bacterial pathogens in which large plasmids play a dominant role in resistance dissemination. In *C. jejuni*, however, limited plasmid diversity should not be interpreted as limited horizontal gene transfer. Rather, the species is highly naturally transformable, and genetic exchange commonly occurs through uptake of exogenous DNA followed by homologous recombination into the chromosome. In this context, the highly open pangenome observed in our dataset supports a model in which adaptation is driven primarily by chromosomal genome plasticity and lineage diversification, facilitated by frequent chromosomal horizontal gene transfer via natural transformation and homologous recombination, rather than by the acquisition of large mobile genetic elements [9,45–47]. The predominance of non-mobilizable genomes therefore indicates a limited large-plasmid burden, while remaining compatible with extensive recombination-mediated evolution.

Taken together, the added contribution of this study is that it places African *C. jejuni* genomic diversity within a global evolutionary framework while simultaneously integrating population structure, pangenome architecture, AMR, virulence, and mobilome features in a single analysis. This synthesis shows that African genomes are not peripheral to global *C. jejuni* diversity; rather, they include globally disseminated lineages, regionally enriched sublineages, and a large accessory gene repertoire that expands the known genomic landscape of the species. Our findings support a unified evolutionary framework in which *C. jejuni* population structure is shaped by three interacting processes: global dissemination of successful generalist lineages, local diversification within regional ecological reservoirs, and continuous genomic plasticity driven by an open pangenome. Within this framework, resistance and virulence follow distinct evolutionary trajectories: AMR is relatively constrained and shaped by regional selective pressures, whereas virulence is highly conserved, reflecting essential biological functions.

This study has important implications for genomic surveillance and public health. First, the pronounced geographic gaps in African genome representation highlight the urgent need for expanded sequencing efforts to achieve equitable global surveillance. Second, the identification of region-specific resistance and virulence patterns suggests that AMR monitoring should incorporate geographic context rather than relying solely on global averages [7–9]. Third, the combination of extensive genomic diversity and conserved functional traits suggests that interventions targeting core biological processes may have broad applicability across lineages.

A key strength of this study is the integration of a large, standardized genome collection with multiple complementary analytical layers, including phylogenomics, pangenome reconstruction, AMR profiling, virulence-factor analysis, and mobilome characterization. The inclusion of African isolates within a broader global comparator dataset allowed regional patterns to be interpreted against the wider *C. jejuni* population structure rather than in isolation. Several limitations should nevertheless be considered. Unlike the East Africa comparator study, which included phenotypic antimicrobial susceptibility testing and directly sampled human and poultry isolates, this analysis relied on publicly available genome data and inferred resistance from genomic determinants. Public repositories are affected by uneven country representation, incomplete host and epidemiologic metadata, inconsistent sampling frames, variable sequencing objectives, and possible overrepresentation of countries or projects with stronger sequencing capacity. Therefore, the observed regional patterns should be interpreted as genomic surveillance signals rather than direct estimates of disease incidence, transmission, or phenotypic resistance prevalence.

Because some public-read assemblies were highly fragmented, estimates of rare gene families and mobilome features may be sensitive to assembly contiguity. These findings should therefore be interpreted cautiously pending confirmation using quality-filtered sensitivity analyses or higher-contiguity genome assemblies. Regional differences may also partly reflect unequal clonal-complex representation across repository datasets.

In conclusion, this study provides a comprehensive genomic framework for understanding *C. jejuni* population structure and functional diversity across Africa and globally. Our results show that *C. jejuni* is characterised by extensive genetic diversity within a globally connected population, a highly dynamic accessory genome, and a conserved functional core. These findings advance our understanding of the evolutionary processes shaping this important foodborne pathogen and provide a foundation for improved genomic surveillance and targeted intervention strategies.

## Data availability

All genome sequence datasets analysed in this study were retrieved from publicly accessible repositories, including the National Center for Biotechnology Information Sequence Read Archive and Assembly resources and corresponding records available through the International Nucleotide Sequence Database Collaboration. Accession identifiers, BioSample records, run accessions, country metadata, host/source information, and derived analytical outputs are provided in Supplementary Data 1.

## Acknowledgements

We acknowledge the investigators and institutions that generated and deposited the *Campylobacter jejuni* genome sequences analyzed in this study. Their public data sharing made this comparative genomic analysis possible.

## Ethical approval and consent to participate

This study analysed publicly available, de-identified microbial genome sequences and associated repository metadata. No new human participants, human biological samples, or live animals were recruited or sampled for this work.

## Author contributions

**Conceptualization:** Abdalah Makaranga, Reuben S. Maghembe, Eliezer Mwakalapa, Pilirani Chisembe, Daud Kassam and Keigo Shibayama. **Methodology, data curation, formal analysis, investigation and visualization:** Samweli Bahati, Abdalah Makaranga and Reuben S. Maghembe. **Writing—original draft preparation:** Samweli Bahati and Abdalah Makaranga. **Writing—review and editing:** Reuben S. Maghembe, Eliezer Mwakalapa, Henry Mung’ong’o, Albino Kalolo, Claus Thomas, Daud Kassam, Pilirani Chisembe and Keigo Shibayama. **Validation and data curation:** Claus Thomas and Henry Mung’ong’o. **Supervision:** Keigo Shibayama, Reuben S. Maghembe and Albino Kalolo. **Project administration:** Reuben S. Maghembe and Keigo Shibayama. All authors reviewed and approved the final version of the manuscript.

## Supplementary material

Four supplementary Excel workbooks accompany this manuscript. Supplementary Data 1 contains genome metadata and accession information; Supplementary Data 2 contains antimicrobial-resistance and virulence-factor profiles; Supplementary Data 3 and Supplementary Data 4 contain plasmid/mobile genetic element analysis outputs and supporting summary tables.

